# A chromosome-level genome assembly of the South African indigenous, Kolbroek pig, *Sus scrofa domesticus*

**DOI:** 10.1101/2025.11.20.689645

**Authors:** Rae Marvin Smith, Annelin Henriehetta Molotsi, Lucky Tendani Nesengani, Thendo Stanley Tshilate, Sinebongo Mdyogolo, Nompilo Lucia Hlongwane, Tracy Masebe, Appolinaire Djikeng, Ntanganedzeni Mapholi

## Abstract

The Kolbroek pig is indigenous to South Africa and a breed of choice for smallholder farmers. This is mainly due to its characteristics, such as disease resistance and adaptability to tropical agroecological environments. Despite these desirable traits, the genomic architecture of this breed has not been explored. In this study, we report a high-quality genome assembly of the South African Kolbroek pig sequenced at 31 x coverage through a combination of PacBio Sequel IIe HiFi and Illumina Novaseq 6000 Omni-C sequencing. The assembled genome resulted in a length of 2.6 Gb in size, including 83 Scaffolds, which consist of 19 chromosome-size scaffolds with 138.7 Mb/48.5 Mb for scaffold/contig N50 respectively. The BUSCO completeness at 95.5 %. Genome annotation and structure prediction identified 22,025 genes with protein-coding potential. The genome provides an opportunity to investigate genetic variation across multiple pig breeds and serves as a genetic resource to develop breeding programs for the conservation and improvement of the Kolbroek pig.

## Background and summary

South Africa is a net importer of pork, importing mainly from other Southern African countries. Commercial breeders dominate the market with a total of 1,450,713 pigs compared to 893,262 for smallholder breeders (BFAP, 2020; DALRRD, 2021). Non-commercial farmers mainly use indigenous breeds, which are characterised by hardiness and tolerance to harsh local environmental conditions ^1^. The Kolbroek is one of these commonly used indigenous pig breeds in South Africa (Figure 1), classified under *Sus scrofa domesticus* (Historically *Sus indica*)^2^. This breed is generally used in smallholder farming systems where it is considered easy to maintain, as it requires low inputs. As with most indigenous breeds, it is suited to the local environment and climate. The Kolbroek pig is characterised by small size and low litter sizes, and therefore exotic breeds are more preferred in the commercial sector. As a result, there have been limited efforts to investigate the genomic architecture of the Kolbroek pig. Previous studies have demonstrated significant genetic diversity among communal indigenous pig breed populations found in South Africa and Zimbabwe, with higher levels of heterozygosity (0.61 - 0.75) than commercial breeds (0.35 - 0.6) ^1^. To demonstrate some level of distinctness and conserved genetic structure of the Kolbroek, Hlongwane et al., ^3^ reported levels of heterozygosity (He = 0.339) and fixation index (*Fst* = 0.468) between Warthog and Kolbroek. However, the authors further show that South African indigenous breeds such as Kolbroek are more inbred and, therefore, would benefit from better resources to enable their conservation.

**Figure 1:**
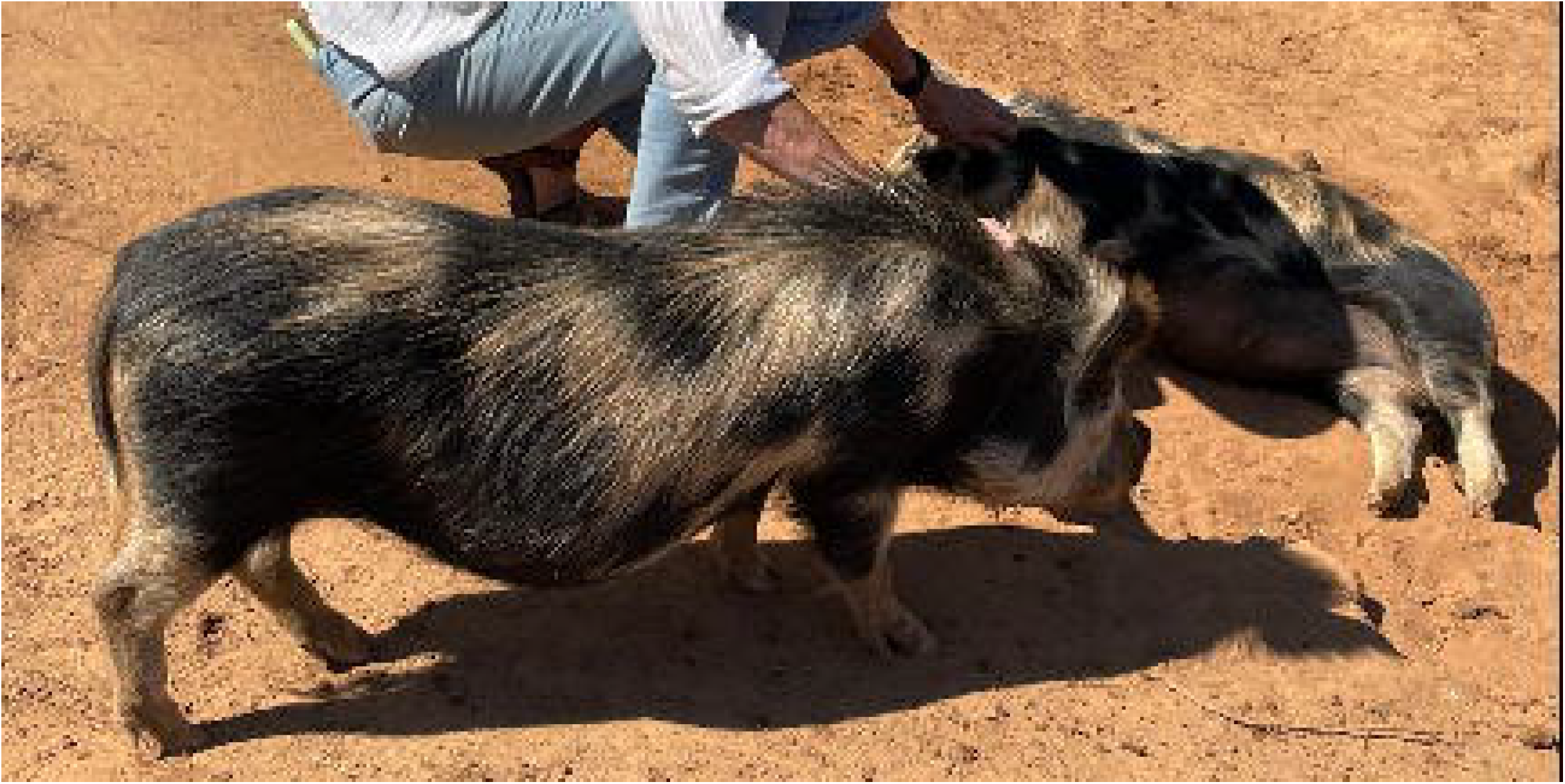
A picture of the Kolbroek Pig.

Economically important traits in pig breeding include fertility, growth, and meat quality. Exotic breeds tend to be preferred over indigenous breeds for commercial pig production purposes ^4^. Regions under natural selection in the Kolbroek genome, such as in chromosome 5, where there are genes associated with motor control, temperature control, control of inflammation, and cell growth, have been identified ^4^. Identification of other useful regions could contribute towards the improvement of these traits. Among other valuable traits, Kolbroek pigs have proven ability to utilise fibrous feeds and tannin-rich red sorghum. They have high parasite tolerance and are suitable for sustainable and organic farmers seeking low input requirements ^1, 5-7^. Research focused on improving traits of economic importance at the genomic level in indigenous pig breeds will be crucial for improving the Kolbroek as a viable commercial option.

Attempts are usually made to harness the desirable qualities of both exotic and indigenous breeds through crossbreeding ^8^. This practice unfortunately leads to the dilution of indigenous breeds and, if not monitored, can eventually lead to the extinction of the breed. To guard against this, there is a need to generate high-quality reference genomes that can be used to develop conservation programs and breeding objectives to improve indigenous breeds. Having a high-quality assembly will enable further investigation of traits such as disease resistance, heat tolerance, adaptability, and the ability to utilize poor-quality forage, as well as facilitate accurate genomic selection. The high-quality genome will assist in accurately identifying quantitative trait loci that are associated with economically important traits and assist in inferring sites of genetic variation ^4^.

Obtaining reference genome assemblies allows geographically representative samples to be incorporated in pangenome graphs, which improves sequence mapping and thus enables single-nucleotide polymorphism (SNP) arrays that are better suited to a particular region. Previous studies have used Illumina Porcine SNP60K and the improved SNP80K, which were developed using commercial breeds and are therefore prone to ascertainment bias when used in indigenous pig breeds. Currently, a major challenge is that genotyping services are still largely unaffordable for farmers who rely on the indigenous breeds that are resilient due to their resistance to local diseases and adaptation to local environmental conditions ^4^. The currently available reference genome of Duroc (*Sscrofa*11.1) is not representative of South African indigenous pig breeds due to selection pressure and population structure ^9^.

In this study, we generated a *de novo* sequenced assembly of a Kolbroek pig, which is a breed that is valued in South Africa by non-commercial farmers. This was done using a combination of 89Gb of high-fidelity (HiFi) sequence reads sequenced at 31 X coverage, and 85Gb of proximity ligated chromatin, (Omni-C) sequence. Workflow developed and curated by the Vertebrate Genome Project, implemented on the Galaxy Bioinformatics platform (Europe), was used. We used a female animal to generate an assembly with a genome size of 2.6 Gb, and contig N50 size of 48.5 Mb. After scaffolding, the assembly was mapped to the 18 autosomes and the X chromosome. The assembly has 18 telomeric regions identified on the chromosomal scaffolds. Repeat elements made up 38.07% of the assembly. The number of protein-coding genes was 22,025. This genome is the first of its kind for the Kolboek breed, and will serve as a resource for the production of specialised SNP panels and comparative genomic analysis against other breeds. This resource will help to identifygenetic variants causing variation in traits of economic importance, which can improve the breed’s value to commercial and non-commercial farmers.

## Methods

### Ethics statement

The procedures for animal handling, sample collection, and all research-related activities were approved by the University of South Africa, Animal Research Ethics Committee (AREC-100818-024), and the Department of Agriculture, Land Reform, and Rural Development (DALRRD) under section 20 of the Animal Diseases Act 1984 (Act 35 of 84) (12/11/1/1/23 (6508 AC)).

### Sample collection and sequencing

A mature pure-breed sow was selected from a Kolbroek pig stud farmer located in the Northwest Province of South Africa. Blood was collected using an EDTA vacutainer vial by a veterinarian, which was immediately placed on dry ice and transported to the laboratory to be stored at -80°C until processing. Genomic DNA was extracted from 200 uL of whole blood using the Nanobind protocol for high molecular weight (HMW) DNA extraction. The HiFi sequencing library was prepared using the SMRTBell® prep kit 3.0 (Pacific Biosciences) following the manufacturer’s instructions and run on the A sequencing library on the Pacific Biosciences (PacBio) Sequel IIe. The Omni-C library preparation was performed from the same sample using the Dovetail Omni-C Kit (Dovetail Genomics), following the manufacturer’s instructions. The resulting Omni-C library was sequenced on the NovaSeq 6000 instrument (Illumina). Initial sequence quality control processing was performed on the SMRTlink Software v11.0 (Pacific Biosciences) as well as FastQC v0.12.1 ^10^.

### Genome assembly

The genome assembly was carried out through a Vertebrate Genome Project workflow ^11^ on the Galaxy platform ^11^. These workflows involve a series of steps that incorporate a selection of bioinformatic tools listed in (Supplementary Table 1). Key steps are to characterize the genome using GenomeScope2 v2.0.1+galaxy0 ^12^. In this step, HiFi reads ^35^ are used to estimate factors such as genome size, heterozygosity, and homozygosity. This step is followed by genome assembly using Hifiasm v0.19.9+galaxy0 ^13^. Hifiasm was performed in Hi-C mode to generate two haplotypes as the kolbroek pig is diploid. These two haplotypes possess structural variants from the paternal and maternal lines. Both assemblies underwent scaffolding with Omni-C reads ^35^, making use of the assemblers, BWA-MEM2 v2.2.1+galaxy1 ^14^ and YAHS v1.2a.2 ^15^. Traces of mitochondrial DNA and other foreign sequences, which may have been incorporated in the assembly, were removed through assembly decontamination, using Kraken 2 v2.1.3+galaxy1 ^16^. This was followed by manual curation using the PretextView software ^17^. K-mer analysis showed an estimated genome size to be 2.6 Gbp (Figure 2). The scaffold N50 was 138.7 Mb, and the contig N50 was 48.5 Mb. Both the clean spectra (CN) plot and the Assembly spectrum (ASM) in supplementary figure 1 display a bimodal distribution with the two peaks at ∼15 and ∼31-fold coverage. The number of contigs for the assembled genomes was 83 scaffolds each. A graphical representation of the assembly statistics for the primary assembly is presented in Supplementary Figure 2.

**Figure 2:**
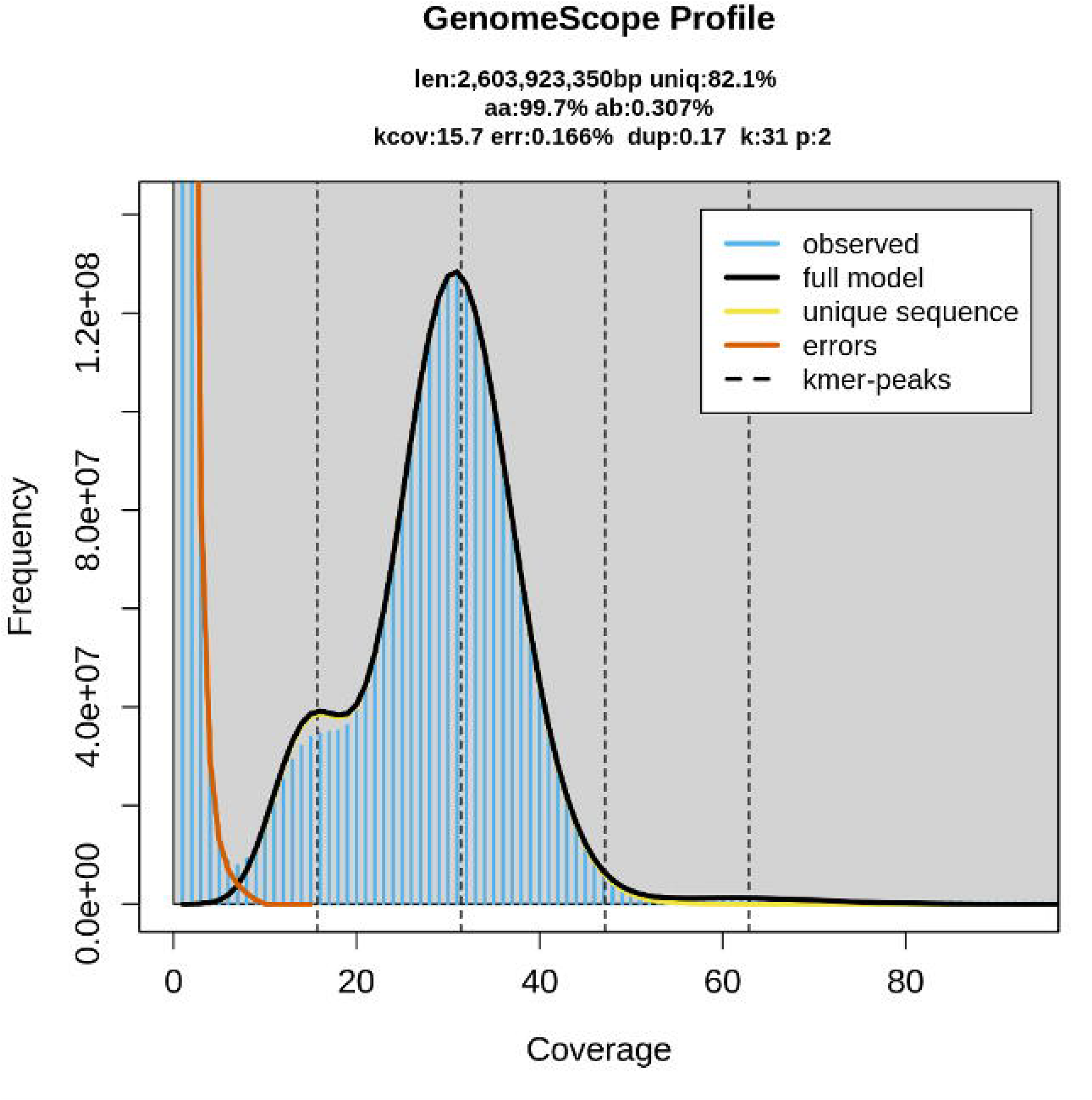
The Genome scope profile generated from HiFi reads. The figure describes the estimated genome size (len), the heterozygosity (ab), and homozygosity (aa), the number of duplicates (dup), the user-specified K-mer size (k), reads that contained errors (err), and the ploidy (p).

### Genome annotation

The transcriptome was not available for support annotation, and thus an *ab initio* approach was used. The genome assembly was soft-masked using RepeatMasker v 4.1.5+galaxy0 ^18^ to mask the repetitive regions in the genome. The repeat elements were then characterised through the modelling software, RepeatModeler v0.09 ^19^, through which RepeatScout v1.0.6 ^20^, RECON v1.5.0 ^20^, and TRF v4.09 ^21^ identified repeat elements that were then annotated using the one_code_to_find_them_all Perl script ^22^. The masked assembly was then used for *de novo* gene annotation using TIBERius v1.1.4 ^23^, which uses a deep-learning *ab initio* gene structure tool for prediction. Structural annotations were determined through filtering the gtf files ^36^ generated through Tiberius and counting the number of predicted elements.

## Data Availability

The completed genome assembly and raw data for the Kolbroek pig were submitted to the National Centre for Biotechnology Information (NCBI) using the Submission number SUB15204155, respectively, with the accession number: JBLUWV000000000 under the Bioproject: PRJNA1227266. Raw reads are available: HIFI reads (SRR32967040), Omni-C reads (SRR32967041), and the Annotation results: (https://doi.org/10.6084/m9.figshare.28754990)

## Technical Validation

The genome assembly was evaluated for completeness, contiguity, correctness, as well as comparisons with related assemblies. Initially, the sequences were quality-checked using FastQC v0.12.1 ^10^. The assemblies were screened using Merqury, which compares K-mers for de novo assembly by ensuring accuracy at the base-level ^24^. Mequry also generates a database file, which is used as an input for genome size estimation through Genomescope 2 v2.0.1+galaxy0 ^12^ and ^25^The Merqury CN plots were generated to check the completeness of the assembly by counting the k-mers. At each stage of the assembly of contigs, purging duplicates, and generating the scaffolds, BUSCO completeness was scored using BUSCO v5.8.0+galaxy1 ^25^. Here, we compared the assembly with the lineage, Cetartiodactyla, which was included in V5 lineage datasets (Figure 3). Also, Gfastats v1.3.11+galaxy0 ^26^ was used to generate statistics at each point of the assembly process, and the final assembly quality statistics were visualized using a snail plot generated through BlobToolKit v 4.0.7+galaxy2 ^27^ (Supplementary Figure 2). The assembly statistics of the Kolbroek genome as compared to other *Sus Scrofa* genomes are provided in Table 1. The genome size and synteny of the Kolbroek are comparable to the NCBI reference genome, Sscrofa11.1 ^33^. Collinearity was assessed using the reference (Figure 4). We identified structural variants that will need to be confirmed through further analysis. This is to distinguish assembly errors from individual/breed structural variation, though the pretext data supports these variants. In particular, translocations in Scaffolds 4, 9, 17, 19 (corresponding to chromosomes 2, 7, 10, and 11 on Sscrofa11.1, respectively). Also, inversions were observed for sequences in two chromosomes, namely, Scaffold 8, 18 and 16 (corresponding with chromosomes 3, and 9 and 10 in Sscrofa11.1, respectively). In terms of telomeric regions, 22 were identified (Supplementary Table 2) using seqtk v1.5+galaxy0 ^28^. Of these, 18 regions are part of the chromosomal scaffolds, and 5 were in the unassembled portion. Repeat elements were masked and characterised (Table 2). Genes were predicted through annotation, and a total of 22,025 genes with a combined coding DNA sequence (CDS) length of 34.23 Mb were identified. CDS length ranged from 201 base pairs at the least to 112,653 base pairs at the largest, with an average of 1,554.3 base pairs. The number of RNA elements, such as small RNA, were 1,235,085. The protein-coding genes ranged from 66 to 37,550 amino acids, and an average protein length of 517.1 amino acids; the corresponding protein sequences obtained from these CDS sequences resulted in 11,388,922 amino acids. Structural features extracted from the annotation file are presented and compared to the reference genome in Table 3. The BUSCO analysis performed to predict gene sequences showed a high-quality annotated genome with 91.5 % completeness, of which 0.5 % were duplicated but complete, and 91.0 %. As a comparison, the reference genome has 98.5 % for the presence of single and duplicate BUSCOs.

**Table 1:** Benchmarking the assembly statistics of the Kolbroek assembly against three public assemblies.

| Statistics | Reference<br>*Sus scrofa*11.1<br><sup>33</sup> | Chenghua <sup>30, 34</sup> | Ningxiang<br><sup>31, 37</sup> | Kolbroek (Current)<br><sup>35</sup> |
| --- | --- | --- | --- | --- |
| Genome size (Gb) | 2.5 | 2.7 | 2.4 | 2.6 |
| Total ungapped length | 2.5 | 2.7 | 2.4 | 2.5 |
| Gaps between scaffolds | 93 | *nd* | *nd* | 77 |
| Number of chromosomes | 20 | 20 | 19* | 19* |
| Number of scaffolds | 705 | 98 | 120 | 77 |
| Scaffold N50 (Mb) | 88.2 | 141.8 | 139 | 138.65 |
| Scaffold L50 | 9 | 8 | 7 | 8 |
| Number of contigs | 1117 | 274 | 421 | 154 |
| Contig N50 (Mb) | 48.2 | 84.7 | 26.1 | 48.5 |
| Contig L50 | 15 | 11 | 29 | 17 |
| GC percent | 42 | 42.5 | 42 | 42 |
| Genome coverage | 65.0 X | 43.0 X | 42.7 X | 31.0X |

nd – no data \* The Y chromosome was not available since the sample was obtained from a female.

**Table 2:** Summary of repeat elements identified in the genome of the Kolbroek pig.

| Repeat element | Number of elements* | Length occupied (Mbp) | Percentage of sequence (%) |
| --- | --- | --- | --- |
| Retroelements | 2,681,508 | 822.9 | 32.43 |
| SINEs: | 1,387,359 | 333.6 | 13.15 |
| LINEs: | 934,163 | 408.2 | 16.09 |
| CRE/SLACS | 0 | 0.0 | 0.00 |
| L2/CR1/Rex | 103,292 | 15.3 | 0.60 |
| RTE/Bov-B | 1378 | 0.2 | 0.01 |
| L1/CIN4 | 829,493 | 392.8 | 15.48 |
| LTR elements: | 359,986 | 81.1 | 3.19 |
| Gypsy/DIRS1 | 0 | 0.0 | 0.00 |
| Retroviral | 343,932 | 79.3 | 3.13 |
| DNA transposons | 207,140 | 33.3 | 1.31 |
| hobo-Activator | 131,861 | 20.6 | 0.81 |
| Tc1-IS630-Pogo | 74,686 | 12.5 | 0.49 |
| MULE-MuDR | 340 | 0.1 | 0.00 |
| PiggyBac | 253 | 0.1 | 0.00 |
| Unclassified: | 34,114 | 69.9 | 2.75 |
| Total interspersed repeats: |  | 926.1 | 36.49 |
| Small RNA: | 1,235,085 | 313.9 | 12.37 |
| Satellites: | 214 | 0.6 | 0.02 |
| Simple repeats: | 677,017 | 34.2 | 1.35 |

\* Most repeats fragmented by insertions or deletions have been counted as one element. Runs of $\geq 20$ X/Ns in query were excluded in % calculations

**Table 3:** Structural Annotations that are observed in the KOLB assembly, which are compared to the reference genome.

| Metric | KOLB<br>(Current)<br>35 | Ref<br>Sscrofa11.1<br>33 |
| --- | --- | --- |
| Number of genes | 22,025 | 27,304 |
| Mean gene length | 31,404.9 | 508,92.6 |
| Number of exons per gene | 8.69371 | 35.5897 |
| Mean exon length (bp) | 178.781 | 331.245 |
| Total genome length (Mb) | 2,109.31 | 13,260.3 |
| Percentage of genes with introns | 81.916 | 0 |
| Mean intron length | 3,879.84 | 0 |
| Total intron length | 657.459 | 0 |
| Average transcript length (bp) | 31,404.9 | 72,301 |

**Figure 3:**
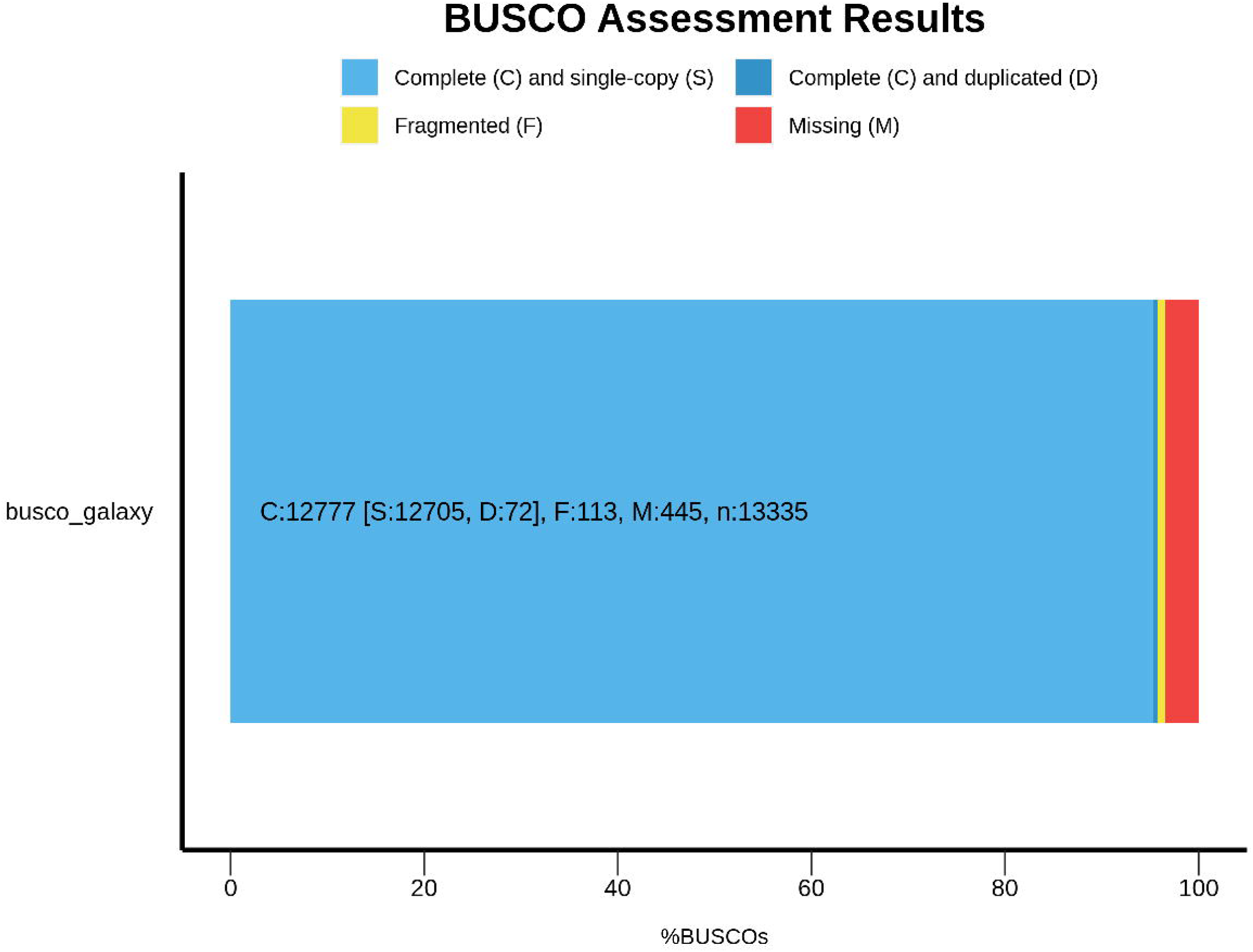
The BUSCO analysis of the primary assembly at the scaffold level. Where the dark and light blue represent the complete single copy and duplicates, respectively. Yellow represents fragmented genes, and red represents what is missing. The assembly was compared to the Cetartiodactyla included in the BUCSO v5 lineage datasets.

**Figure 4:**
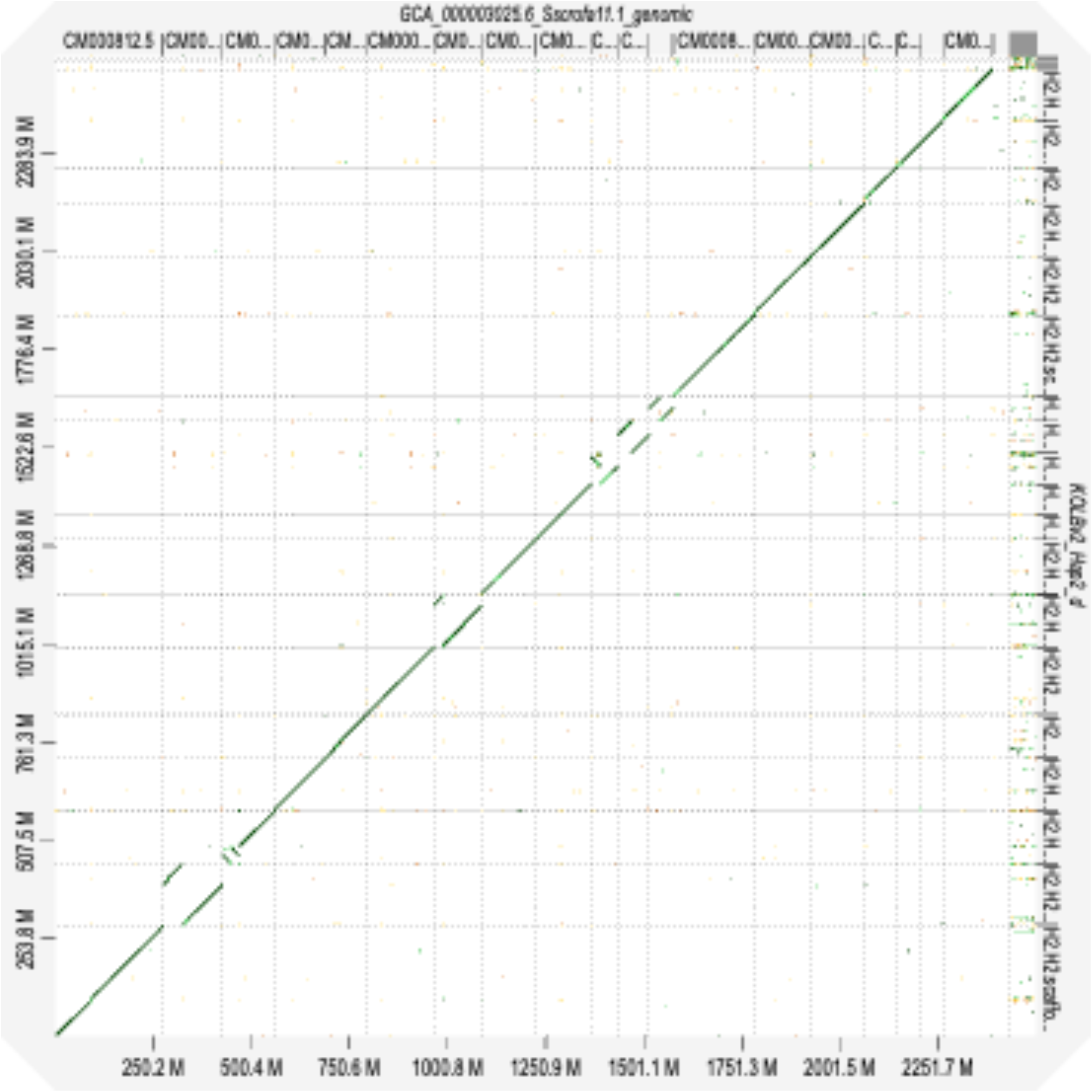
Collinearity plot showing the scaffolds that match with the reference genome, Sscrofa11.1, and the Kolbroek assembly generated in D-genies ^29^.

## Code availability

Genetic analysis was performed on the Galaxy Europe platform (https://usegalaxy.eu) and the workflow that was used is on the Vertebrate Genome Project (VGP) pipeline (https://galaxyproject.org/projects/vgp/workflows/ ). The tools and their versions associated with the VGP assembly pipeline are listed in Supplementary Table 1. Moreover, additional analyses that were performed are also listed. This pipeline is under development; thus, the versions that were used are specified. Where incompatibility issues were encountered, alternate versions were used, which provided options for the required input data. The VGP group updates its pipelines to incorporate newer versions of software or resolve dependency issues.

## NCBI Accession number

BioProject: PRJNA1227266. Accession Sus scrofa (JBLUWV000000000 ; BioSample SAMN46977218)

## Authors’ contributions

Conceptualization: RMS, LTN, NOM, SM, AD; Data Curation: RMS, AHM; Formal Analysis: RMS, LTN, TT, SM, AHM; Funding Acquisition: NOM, TM; Investigation: RMS; Methodology: RMS, LTN, SM, AH; Project Administration: NOM, TM; Resources: LTN, SM, Software: RMS, AHM, TT; Validation: AHM, RMS, LTN, SM, TT; Visualisation: RMS; Writing of the original draft: RMS, AHM; Reviewing and editing of the manuscript: All authors.

## Acknowledgments

We acknowledge funding and support from the University of South Africa for funding In addition, we acknowledge the Vertebrate Genome Project for their support when executing the pipeline. We would also like to thank the Staff at Inqaba Biotech for completing the library preparation of the collected samples and as well as completing the HiFi Sequencing. We would like to thank the staff at the University of Stellenbosch, where the Omni-C data was produced. We would like to acknowledge Prof Jasper Rees, Dr Sikhumbuzo Mbizeni, and Dr Thivhilaheli Richard Netshirovha for their assistance during sample collection and analyses. We would also like to that Prof Cuthbert Banga for his assistance with critical reading. We would like to acknowledge Galaxy Europe for hosting the data and supplying computing resources. This article forms part of the objectives for the Africa Biogenome Project. A special thanks to the team in Mapholi Labs for the resources and for managing the project.

