## Supplementary material for "A chromosome-level genome assembly of the South African indigenous, Kolbroek pig, *Sus scrofa domesticus*": Supplimentary Data and figures

**Supplementary information**

**Supplementary Table 1**: A summary of the assembly pipeline that was used, along with versions available on https://galaxy.eu. The table also indicates which VGP workflow and version where available, was used to track galaxy-associated information.

| **VGP1** | **v0.1.7** | | |
|---|---|---|---|
| step | Tool | version | Reference |
| **K-mer analysis** | meryl | 1.3+galaxy6 | 1 |
| | genomescope | 2.0.1+galaxy0 | 2 |
| **VGP 4** | **V0.2.2** | | |
| step | Tool | version | Reference |
| **Contig assembly** | cutadapt | 4.9+galaxy1 | 15 |
| | multiqc | 1.11+galaxy1 | 16 |
| | hifiasm | 0.19.9+galaxy0 | 3 |
| | gfastats | 1.3.6+galaxy0 | 4 |
| | ggplot2_point | 3.4.0+galaxy1 | 5 |
| | busco | 5.5.0+galaxy0 | 6 |
| | merqury | 1.3+galaxy4 | 13 |
| **VGP6** | **v0.4** | | |
| step | Tool | version | Reference |
| **Purge Duplicates** | minimap2 | 2.28+galaxy0 | |
| | purge_dups | 1.2.6+galaxy0 | https://github.com/dfguan/purge_dups |
| | busco | 5.5.0+galaxy0 | |
| | gfastats | 1.3.6+galaxy0 | |
| | merqury | 1.3+galaxy4 | |
| | ggplot2_point | 3.4.0+galaxy1 | |
| **VGP8** | **v0.2.8** | | |
| **step** | **Tool** | **version** | Reference |
| **Scaffold assembly** | gfastats | 1.3.6+galaxy0 | |
| | bwa_mem2 | 2.2.1+galaxy1 | 7 |
| | bellerophon | 1.0+galaxy1 | 8 |
| | pretext_map | 0.1.9+galaxy1 | https://github.com/sanger-tol/PretextMap |
| | YaHS | 1.2a.2+galaxy2 | 9 |
| | pretext_snapshot | 0.0.3+galaxy2 | https://github.com/sanger-tol/PretextSnapshot |
| | busco | 5.5.0+galaxy0 | |
| | datamash | 1.8+galaxy0 | |
| | bedtools_bamtobed | 2.31.1+galaxy0 | 24 |
| | ggplot2_point | 3.4.0+galaxy1 | |

| | | | |
|---|---|---|---|
| **VGP9** | **v0.3** | | |
| **step** | **Tool** | **version** | |
| **Decontamination** | kraken2 | 2.1.3+galaxy1 | [10] |
| | gfastats | 1.3.6+galaxy0 | |
| **step** | **Tool** | **version** | |
| Colinearity | D-genies* | | [17] |
| Summary | Blobtools | 4.0.7+galaxy2 | [18] |
| Summary | BlobToolKit | 4.0.7+galaxy2 | |
| Manual Curation | PretextView* | 0.04 | https://github.com/sanger-tol/PretextView |
| | YaHS | 1.2a.2+galaxy2 | |
| Annotation | RepeatMasker | 4.1.5+galaxy0 | |
| Annotation | Repeat Modeler | 2.0.5+galaxy0 | |
| Annotation | RepeatScout | v1.0.6 | [11] |
| Annotation | RECON | v1.5.0 | [11] |
| Annotation | TRF | v4.09 | [12] |
| Annotation | Tiberius* | v1.1.4 | [13] |
| Quality control | FastQC | 0.12.1 | [11] |
| Mitogenome assembly | MitoHiFi | 3 | [14] |

*not available on Galaxy at the time of analysis

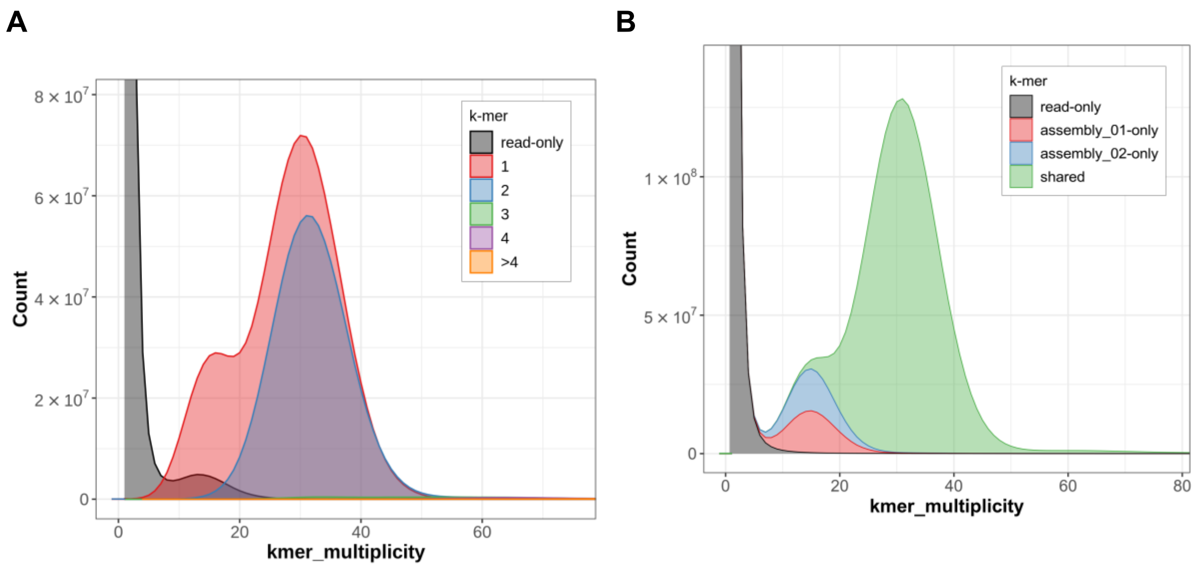

**Supplementary Figure 1**: A CN plot (A) that was generated for the hap 1 assembly showing the level of K-mer multiplicity for HiFi reads and an Assembly spectrum (ASM) plot (B) showing the K-mers identified, which are unique for each assembly at the contig level, as well as those that are shared.

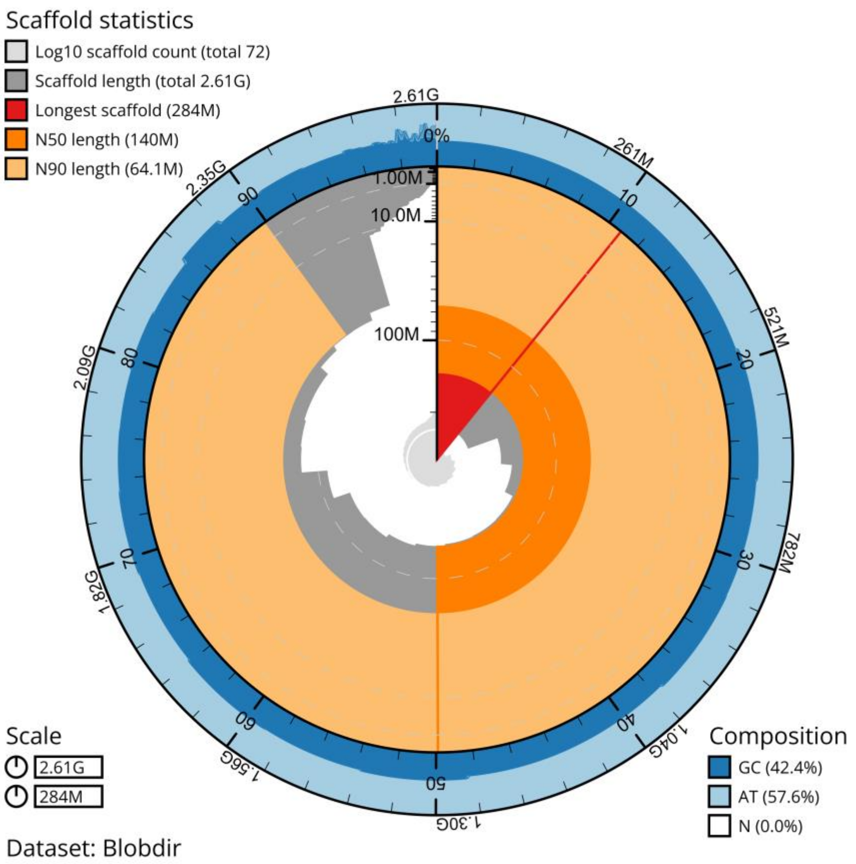

**Supplementary Figure 2**. A snail plot that graphically shows the quality of the primary genome assembly. The plot of the circumference of the circle represents the size (bp) of the genome, with the inner spiral describing the cumulative level of representation of each of the scaffolds. Dark red represents the first and longest scaffold, dark orange shows the scaffold N50 value, and light orange shows the included data for scaffold N90. The dark and light blue values show the percent GC and AT content respectively[15].

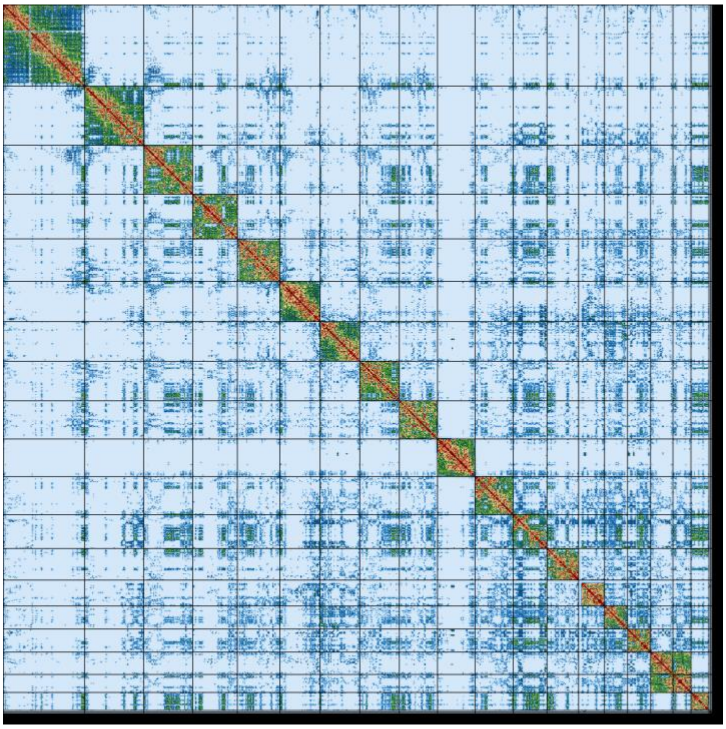

**Supplementary Figure 3:** An Omni-C contact map generated against the Kolbroek primary assembly was generated, showing the chromosome-level scaffolds. The deeper intensity colour from light blue to dark red indicates the strength of the interactions between Omni-C reads and the assembly sequences. The scaffolds were ordered by size.

**Supplementary Table 2**: The results from the telomere analysis using the seqtk_telo application. The table displays the start and end regions, telomere lengths, as well as the average length for those that are identified. The bold Scaffold names indicate those that aligned to the reference during the Collinearity plot.

| Chromosome | Start | End | Telomere length |
|---|---|---|---|
| **scaffold_01** | 0 | 10249 | 10249 |
| **scaffold_01** | 284068673 | 284075074 | 6401 |
| **scaffold_02** | 0 | 10432 | 10432 |
| **scaffold_02** | 222666662 | 222672346 | 5684 |
| **scaffold_05** | 0 | 3774 | 3774 |
| **scaffold_06** | 142879150 | 142883766 | 4616 |
| **scaffold_07** | 0 | 4990 | 4990 |
| **scaffold_07** | 142073233 | 142083538 | 10305 |
| **scaffold_08** | 139841255 | 139850298 | 9043 |
| **scaffold_09** | 0 | 3970 | 3970 |
| **scaffold_09** | 136850103 | 136854792 | 4689 |
| **scaffold_11** | 0 | 12397 | 12397 |
| **scaffold_12** | 0 | 3090 | 3090 |
| **scaffold_14** | 80007280 | 80009781 | 2501 |
| **scaffold_17** | 0 | 803 | 803 |
| **scaffold_17** | 68683873 | 68687960 | 4087 |
| **scaffold_19** | 0 | 1561 | 1561 |
| **scaffold_20** | 0 | 6744 | 6744 |
| scaffold_23 | 0 | 5250 | 5250 |
| scaffold_27 | 6018641 | 6020598 | 1957 |
| scaffold_33 | 0 | 2545 | 2545 |
| scaffold_35 | 0 | 4536 | 4536 |
| scaffold_72 | 18730 | 28028 | 9298 |
| Mean length | | | 5605.30 |
